# diverse-seq: an application for alignment-free selecting and clustering biological sequences

**DOI:** 10.1101/2024.11.10.622877

**Authors:** Gavin Huttley, Katherine Caley, Robert McArthur

## Abstract

The algorithms required for phylogenetics — multiple sequence alignment and phylogeny estimation — are both compute intensive. As the size of DNA sequence datasets continues to increase, there is a need for a tool that can effectively lessen the computational burden associated with this widely used analysis.

`diverse-seq` implements computationally efficient alignment-free algorithms that enable efficient prototyping for phylogenetic workflows. It can accelerate parameter selection searches for sequence alignment and phylogeny estimation by identifying a subset of sequences that are representative of the diversity in a collection. We show that selecting representative sequences with an entropy measure of *k*-mer frequencies correspond well to sampling via conventional genetic distances. The computational performance is linear with respect to the number of sequences and can be run in parallel. Applied to a collection of 10.5k whole microbial genomes on a laptop took ∼8 minutes to prepare the data and 4 minutes to select 100 representatives. `diverse-seq` can further boost the performance of phylogenetic estimation by providing a seed phylogeny that can be further refined by a more sophisticated algorithm. For ∼1k whole microbial genomes on a laptop, it takes ∼1.8 minutes to estimate a bifurcating tree from mash distances.

The `diverse-seq` algorithms are not limited to homologous sequences. As such, they can improve the performance of other workflows. For instance, machine learning projects that involve non-homologous sequences can benefit as representative sampling can mitigate biases from imbalanced groups.

`diverse-seq` is a BSD-3 licensed Python package that provides both a command-line interface and `cogent3` plugins. The latter simplifies integration by users into their own analyses. It is available via the Python Package Index and GitHub.

**Statement of need:** Accurately selecting a representative subset of biological sequences can improve the statistical accuracy and computational performance of data sampling workflows. In many cases, the reliability of such analyses is contingent on the sample capturing the full diversity of the original collection (e.g. estimating large phylogenies Parks et al., 2018; Zhu et al., 2019). Additionally, the computation time of algorithms reliant on numerical optimisation, such as phylogenetic estimation, can be markedly reduced by having a good initial estimate.

Existing tools to the data sampling problem require input data in formats that themselves can be computationally costly to acquire. For instance, tree-based sequence selection procedures can be efficient, but they rely on a phylogenetic tree or a pairwise genetic distance matrix, both of which require alignment of homologous sequences (Balaban et al., 2019; e.g. Widmann et al., 2006). Adding both the time for sequence alignment and tree estimation presents a barrier to their use.

The `diverse-seq` sequence selection algorithms are linear in time for the number of sequences and more flexible than published approaches. While the algorithms do not require sequences to be homologous, when applied to homologous sequences, the set selected is comparable to what would be expected based on genetic distance. The `diverse-seq` clustering algorithm is linear in time for the combined sequence length. For homologous sequences, the estimated trees are approximations to that estimated from an alignment by IQ-TREE2 (Minh et al., 2020).

## Definitions

A *k*-mer is a subsequence of length *k*, and the frequency of each *k*-mer in a sequence forms a *k*-mer probability vector. The Shannon entropy is a measure of “uncertainty” in a probability vector and is commonly used in sequence analysis (e.g. Schneider & Stephens, 1990). The Shannon entropy is calculated as *H* = − ∑_*i*_ *p*_*i*_ log_2_ *p*_*i*_ where *p*_*i*_ is the probability of the *i*-th *k*-mer. As an indication of the interpretability of Shannon entropy, a DNA sequence with equifrequent nucleotides has the maximum possible *H* = 2 (high uncertainty) while a sequence with a single nucleotide has *H* = 0 (no uncertainty).

Shannon entropy is integral to other statistical measures that quantify uncertainty (Lin, 1991), including Jensen-Shannon divergence (JSD), which we employ in this work. As illustrated in Table 1, the magnitude of JSD reflects the level of relatedness amongst sequences via the similarity between their *k*-mer probability vectors. For a collection of *N* DNA sequences 𝕊, define *f*_*i*_ as the *k*-mer frequency vector for sequence *s*_*i*_. The JSD for the resulting set of vectors, 𝔽, is

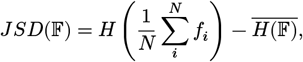

where the first term corresponds t o the Shannon entropy o f the mean o f t he *N* probability vectors and the second term 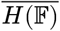 is the mean of their corresponding Shannon entropies. For vector *f*_*i*_ ∈ 𝔽 its contribution to the total JSD of 𝔽 is

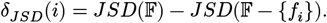

**Table 1.**
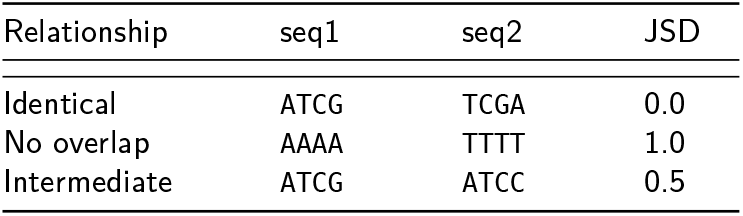
Jensen-Shanon Divergence (JSD) for different relationships between two sequences with *k* = 1.

From the equation, it is apparent that to update the JSD of a collection efficiently, we need only track *k*-mer counts, total Shannon entropy and the number of sequences. Thus, the algorithm can be implemented with a single pass through the data.

To facilitate the description below, we define the record with the minimum δ _*JSD*_ as

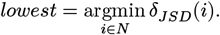

## Algorithms

### *dvs* *subcommand:* *prep* *converts sequences into* *numpy* *arrays for faster processing*

The `prep` sub-command converts plain text sequence data into an on-disk storage format for efficient access in the other steps. A user can provide either fasta or GenBank formatted DNA sequence files. The sequences are converted into unsigned 8-bit integer `numpy` arrays and stored in a single HDF5 file on disk. The resulting `.dvseqs` file is required for all the sub-commands.

## Selection of representative sequences

### *dvs* *subcommands**nmost* *samples the n sequences that increase JSD most;* *max* *samples sequences that maximise a user specified statistic, either the standard deviation or the coefficient of variation of* δ_*JSD*_

The following optimisations have been employed to make the algorithm for computing the JSD scalable in terms of the number of sequences.

1. Sequence data is BLOSC2 compressed as unsigned-8 bit integers and saved in HDF5 format on disk.
2. `numba`, a just-in-time compiler, is used for the core algorithms producing *k*-mers and their counts, providing a significant speed up over a pure python implementation (Lam et al., 2015).
3. Sequence loading and *k*-mer counting is triggered when a sequence record is considered for inclusion in the divergent set, reducing the memory required to that for the user-nominated size.

The `nmost` algorithm defines an exact number of sequences to be selected that maximise the JSD. The order of input sequences is randomised and the selected set is initialised with the first *n* sequences. As shown in Figure 1, for each of the remaining sequences, if adding it to the set 𝔽 − {*lowest*} increases JSD, it replaces *lowest*. The `max` algorithm differs from `nmost` by defining lower and upper bounds for the number of sequences in the divergent set. It further amends the within-loop condition (Figure 2), allowing the number of sequences in the set to change when a statistical measure of δ_*JSD*_ variance increases. We provide users a choice of two measures of variance in δ_*JSD*_: the standard deviation or the coefficient of variation.

**Figure 1.**
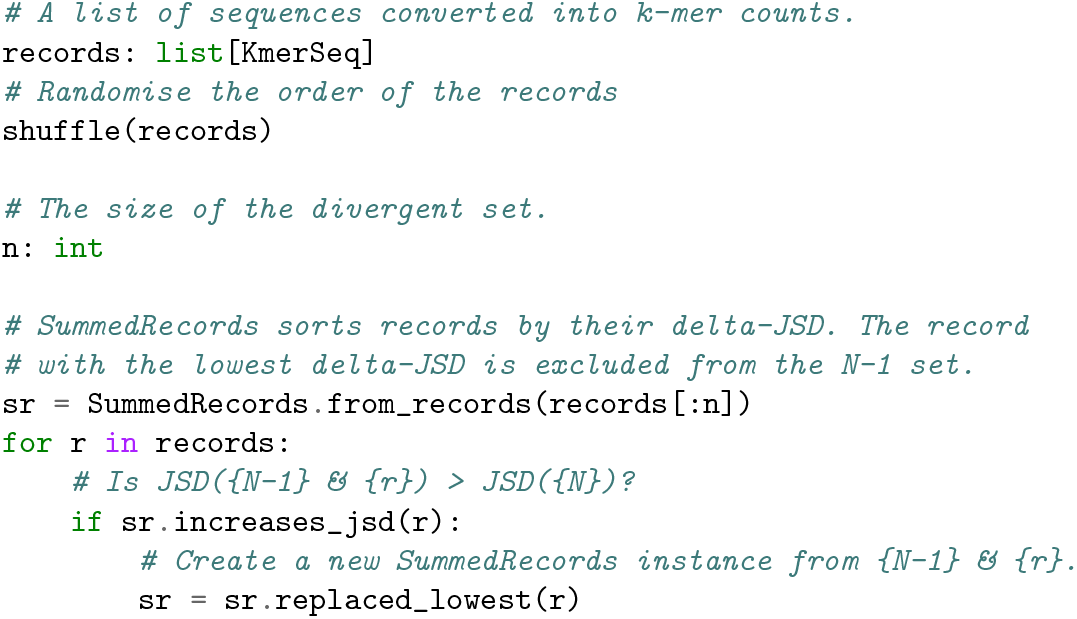
The `diverse-seq nmost` algorithm.

**Figure 2.**
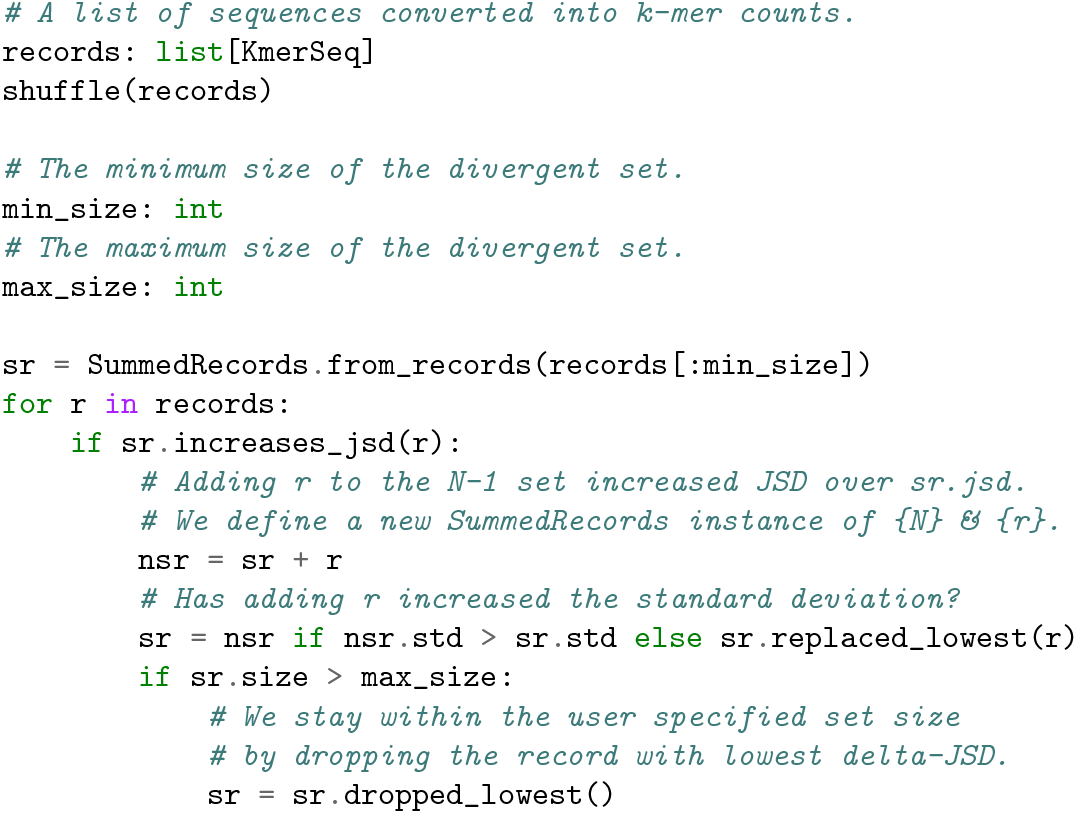
The `diverse-seq max` algorithm. This includes upper and lower bounds for the size of the divergent set and amends the within-loop condition of `nmost`. The set size is increased when a record that increases JSD also increases the standard deviation of δ_*JSD*_.

## Constructing a tree from *k*-mers

### *dvs* *subcommand:* *ctree* *estimates a phylogenetic tree from unaligned sequences using mash distances*

The mash distance (Ondov et al., 2016) estimates the proportion of changes between two sequences and can be computed in near linear time. It approximates the Jaccard distance between the *k*-mer sets of the two sequences (the proportion of shared *k*-mers) from a subset of all *k*-mers in the two sequences. This subset, the MinHash sketch, is the *sketch size* smallest *k*-mers when sorted by a hash function. We apply agglomerative clustering with average linkage (Murtagh & Contreras, 2012) to the pairwise mash distances to estimate a phylogenetic tree from unaligned sequences. We allow the user to select the distance metric, *k* and the sketch size.

### dvs cogent3 apps

We provide `dvs_nmost`, `dvs_max`, `dvs_ctree` and `dvs_par_ctree` as cogent3 apps. For users with cogent3 installed, these are available at runtime via the cogent3 function `get_app()`. The apps mirror the settings from their command-line implementation but differ in that they operate directly on a sequence collection, skipping conversion to disk storage. The `dvs_nmost` and `dvs_max` directly return the selected subset of sequences, `dvs_ctree` and `dvs_par_ctree` returns the estimated phylogenetic tree. The `dvs_par_ctree` app runs in parallel on a single alignment. The `dvs_max` and `dvs_ctree` apps are demonstrated in the `plugin_demo.ipynb` notebook.

## Performance

### Selection of representative sequences

#### Recovery of representatives from synthetic knowns

We evaluate the ability of `dvs max` to recover known divergent lineages using simulated data. The experimental design was intended to assess whether rare sequences can be recovered. We defined four distinct sequence compositions and two distinct “pool” compositions: *balanced*, in which each sequence family was present at equal frequency, or *imbalanced*, where one sequence occurred at 1%, another 49% and the remainder at 25% each. In each scenario, we simulated a total of 50 sequences. If `dvs max` identifies a set of precisely 4 sequences with one pool representative, this is counted as a success. As shown in Figure 3, the primary determinant of the success was the length of the simulated sequences.

**Figure 3.**
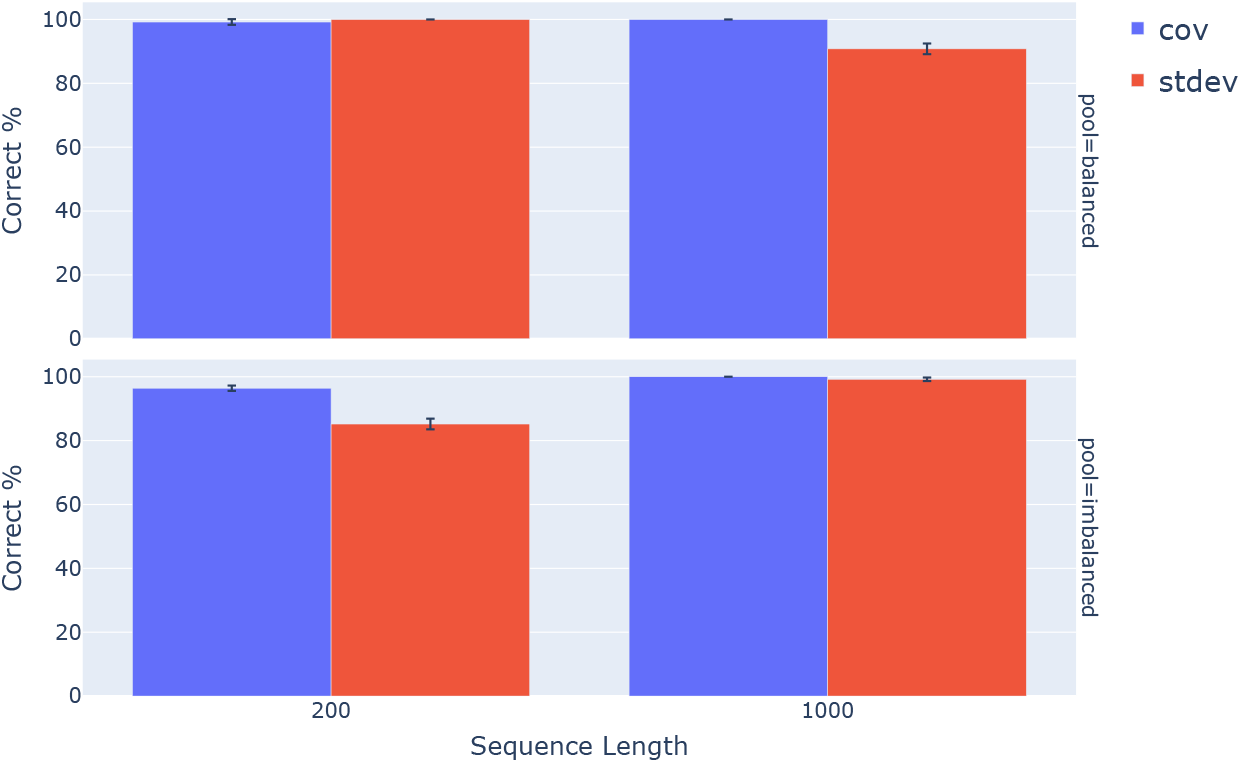
Identification of representatives of known groups is affected by sequence length. `dvs max` identified representatives of known groups in both *balanced*, and *imbalanced* pools.

#### The selected sequences are phylogenetically diverse

For homologous DNA sequences, increasing the amount of elapsed time since they shared a common ancestor increases their genetic distance due to time-dependent accumulation of sequence changes. We expect that the JSD between two sequences will also increase proportional to the time since they last shared a common ancestor. We therefore pose the null hypothesis that if JSD is not informative, then the minimum pairwise genetic distance amongst *n* sequences chosen by `diverse_seq` will be approximately equal to the minimum pairwise genetic distance between a random selection of *n* sequences. Under the alternate hypothesis that JSD is informative, the minimum genetic distance between sequences chosen by `diverse_seq` will be larger than between randomly selected sequences. We test this hypothesis using a resampling statistic (Sokal & Rohlf, 1995, p. 808), estimating the probability of the algorithmic choice being consistent with the null hypothesis. This probability is calculated as the proportion of 1000 randomly selected sets of sequences whose minimum genetic distance was greater or equal to that obtained from the sequences chosen by `dvs max`. We further summarised the performance of the `dvs` commands as the percentage of loci which gave a *p*-value less than 0.05. A bigger percentage is better.

We addressed the above hypothesis using 106 alignments of protein coding DNA sequences from the following 31 mammals: Alpaca, Armadillo, Bushbaby, Cat, Chimp, Cow, Dog, Dolphin, Elephant, Gorilla, Hedgehog, Horse, Human, Hyrax, Macaque, Marmoset, Megabat, Microbat, Mouse, Orangutan, Pig, Pika, Platypus, Rabbit, Rat, Shrew, Sloth, Squirrel, Tarsier, Tenrec and Wallaby. The sequences were obtained from Ensembl.org (Harrison et al., 2024) and aligned using cogent3’s codon aligner (Knight et al., 2007). The genetic distance between the sequences was calculated using the paralinear distance (Lake, 1994).

The results of the analysis (Figure 4) indicated the sucess of `dvs max` in identifying genetically diverse sequences was principally sensitive to the choice of *k*. While Figure 4(a) showed close equivalence between the statistics, Figure 4(b) indicates the size of the selected set using the standard deviation was systematically lower than for the coefficient of variation. The result from the `dvs nmost` analysis, which performed using the minimum set size argument given to `dvs max` is represented by the *JSD*(𝔽) statistic.

**Figure 4.**
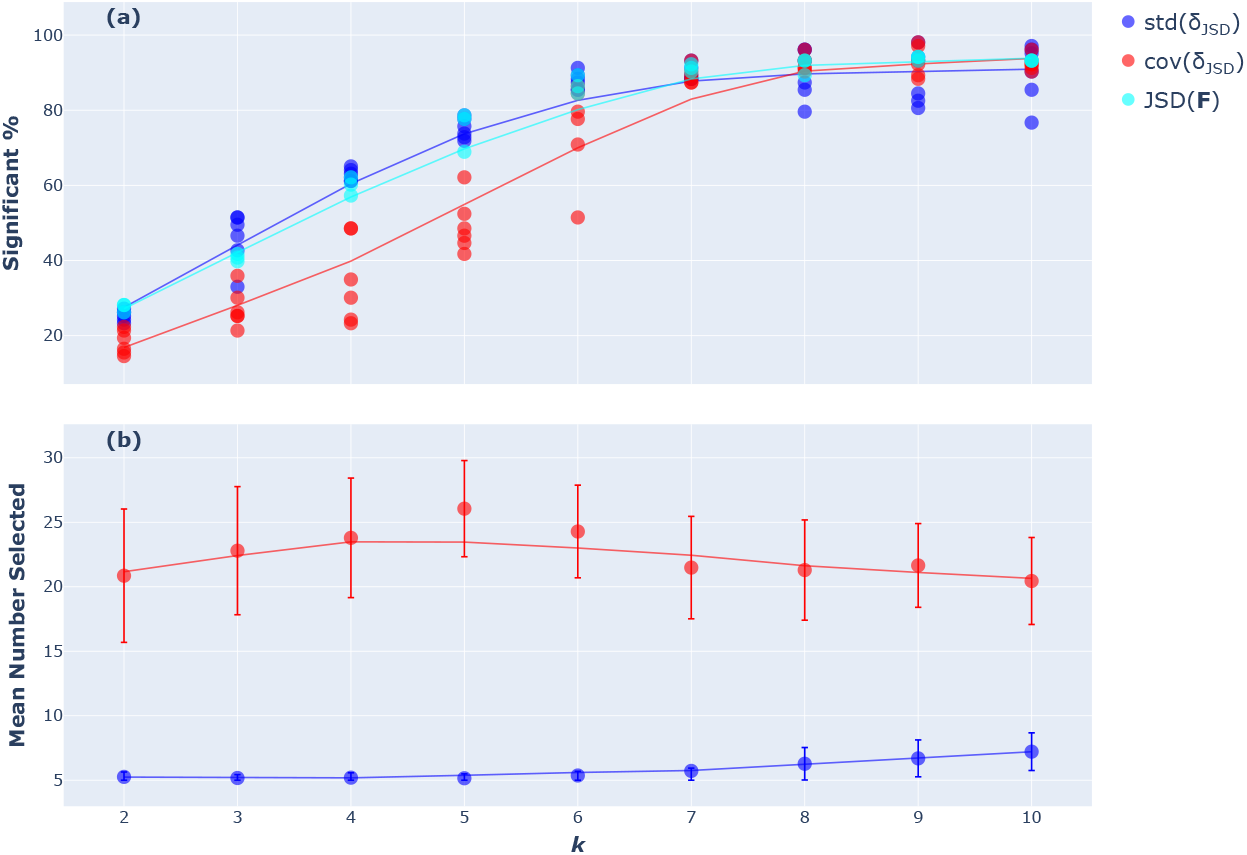
The statistical performance of `dvs max` in recovering representative sequences is a function of *k* and the chosen statistic. The minimum and maximum allowed set sizes were 5 and 30, respectively. `dvs nmost` is represented by *JSD*(𝔽) run with n=5. Trendlines were estimated using LOWESS (Cleveland, 1979). (a) *Significant %* is the percentage of cases where `dvs max` was significantly better (*p* − value ≤ 0.05) at selecting divergent sequences than a random selection process. (b) The mean and standard deviations of the number of sequences selected by `dvs max`.

**Figure 5.**
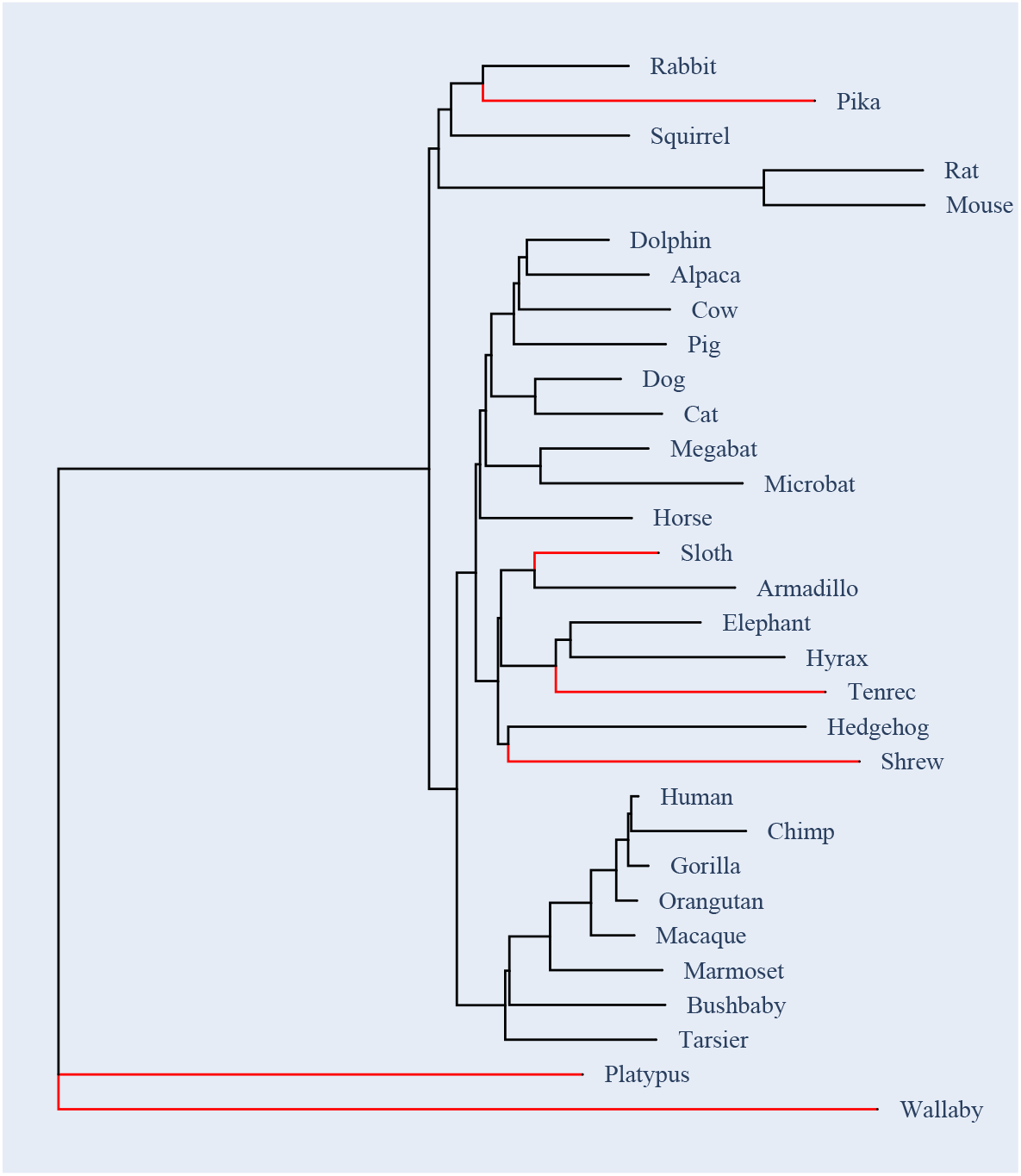
Result of applying the `dvs_max` app to a single sequence alignment. The phylogenetic tree was estimated using Neighbour-Joining (Saitou & Nei, 1987) from the pairwise paralinear distances (Lake, 1994). The branch to the sequence selected by `dvs_max` are shown in red. See the `plugin_demo` notebook for the code used to produce this figure.

#### Computational performance

As shown in Figure 6, the compute time was linear with respect to the number of sequences, shown using random samples of microbial genomes from the 960 REFSOIL dataset (Choi et al., 2017). We further trialled the algorithm on the dataset of Zhu et al. (2019), which consists of 10,560 whole microbial genomes. Using 10 cores on a MacBook Pro M2 Max, application of `dvs prep` followed by `dvs nmost` took 8’9” and 3’45” (to select 100 sequences) respectively. The RAM memory per process was ∼300MB.

**Figure 6.**
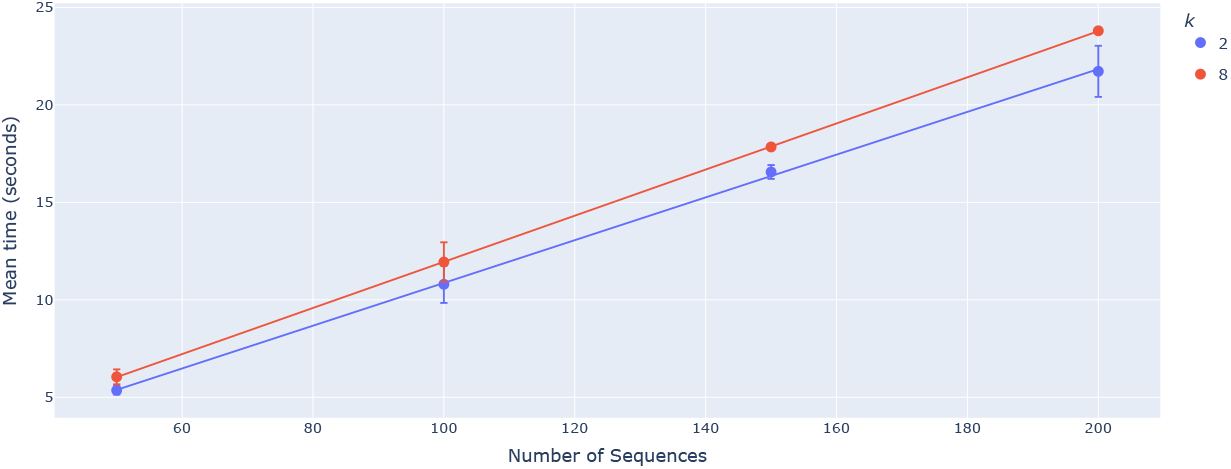
`dvs max` exhibits linear time performance with respect to the number of microbial genome sequences. Three replicates were performed for each condition. For each repeat, sequences were randomly sampled without replacement from the 960 REFSOIL microbial dataset (Choi et al., 2017).

### Constructing trees from *k*-mers

We use the mammals dataset to evaluate the statistical performance of the `ctree` method. All sequences were concatenated and a phylogenetic tree was estimated from this alignment with different *k*-mer sizes and sketch sizes. The trees generated by `dvs ctree` are compared to the maximum likelihood tree found by IQ-TREE2 (Minh et al., 2020) using a general time-reversible model (Tavaré, 1986) on the concatenated alignment.

The likelihood of the generated trees changes with *k* (Figure 7). When *k* ≤ 5, the *k*-mers are non-unique (all mash-distances are zero) and the method generates a caterpillar tree. It is not until *k* = 8 when the caterpillar tree is statistically outperformed. As *k* increases further, the likelihood approaches that of IQ-TREE2 but plateaus before it at *k* = 12. Figure 8 shows how the likelihood of the generated trees changes as the sketch size increases for varying *k* ≥ 8.

**Figure 7.**
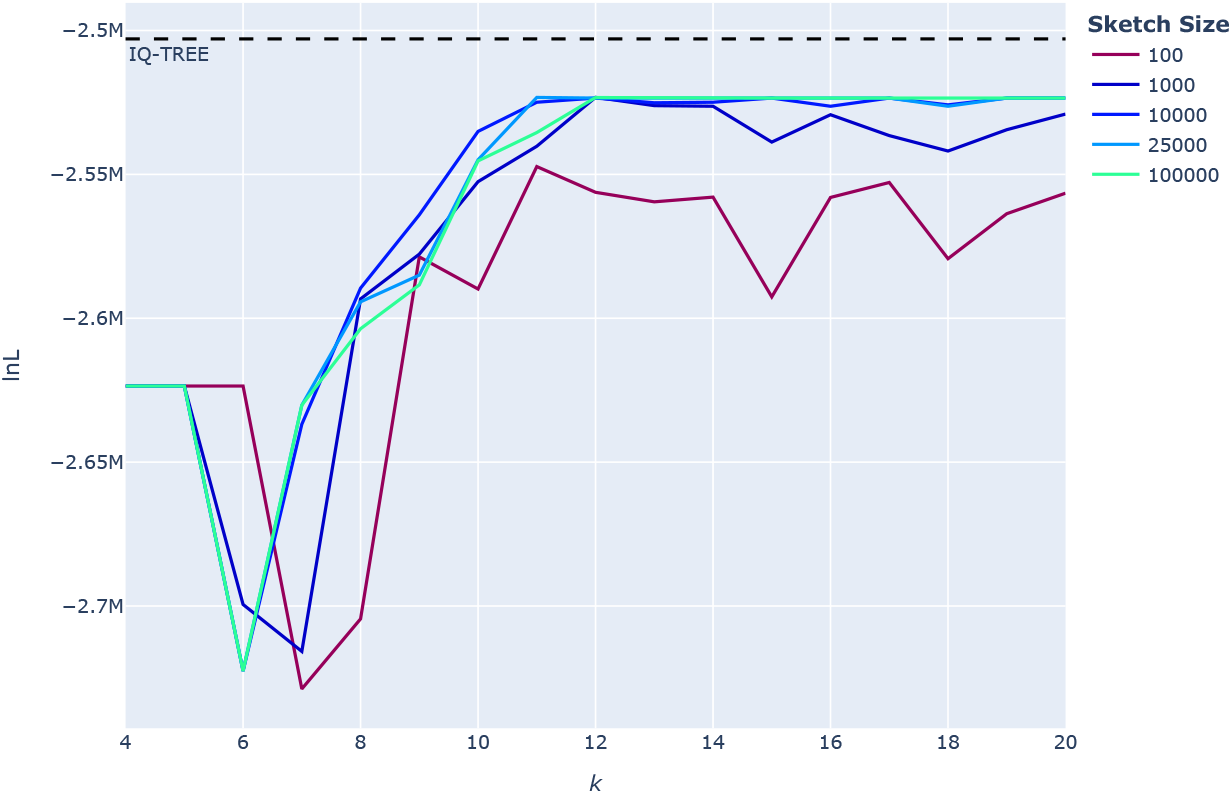
Statistical performance of the `dvs_ctree` app on the concatenated mammals alignment as the *k*-mer size increases. The likelihood of trees, represented as the log-likelihood (lnL), generated by the app is compared to the maximum likelihood tree found by IQ-TREE2 (Minh et al., 2020). For large enough sketch sizes, the likelihood approaches that of IQ-TREE2 and plateaus beyond a *k*-mer size of ∼12.

**Figure 8.**
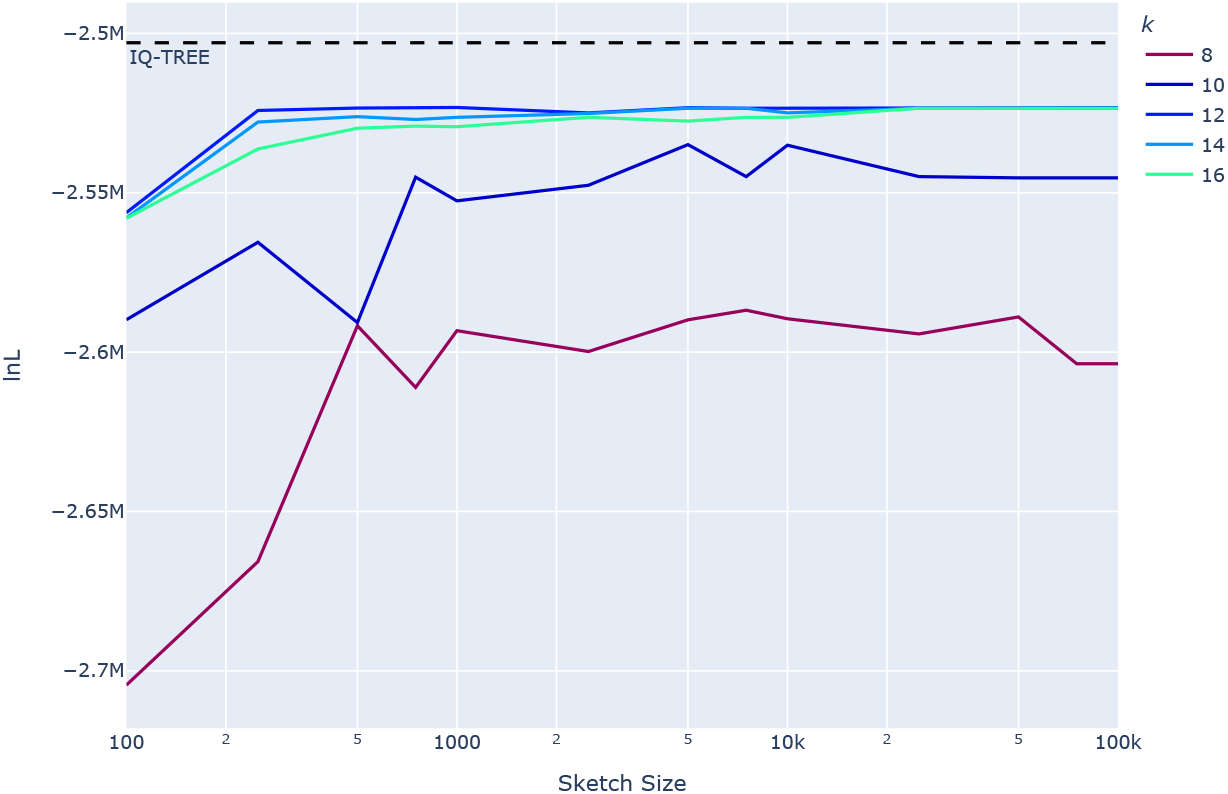
Statistical performance of the `dvs_ctree` app on the concatenated mammals alignment as the sketch size increases. The likelihood of trees, represented as the log-likelihood (lnL), generated by the app is compared to the maximum likelihood tree found by IQ-TREE2 (Minh et al., 2020). For optimal *k*-mer sizes, the likelihood approaches that of IQ-TREE2 and plateaus beyond a sketch size of ∼2500.

Sketch size also impacts on the tree likelihood. An upward trend with increasing sketch size was evident with the likelihood approaching but not reaching that found by IQ-TREE2 on the aligned data. For the best performing values of *k*, there was no benefit to increasing sketch size beyond ∼2,500.

#### Computational performance

While the time complexity of the standard algorithm for agglomerative clustering is 𝒪(*n*^3^), with respect to the number of sequences, the number of sequences is often small in comparison to the sequence lengths. As such, it was found that the most time consuming step of the algorithm was within the distance calculation. The pairwise distance calculation is done in two steps. First, is the constructing the MinHash sketch for each sequence (a subset of sketch size *k*-mers), followed by the computation of distances from these sketches (Ondov et al., 2016). The expected runtime for constructing the MinHash sketch for a sequence is 𝒪(*l* +*s* log *s* log *l*) where *l* is the length of the sequence, and *s* is the sketch size. This is linear with respect to *l* when *l* >> *s*. Hence, the time complexity for constructing all MinHash sketches is linear with respect to the combined length of all sequences. The time complexity for calculating the distance between two sequences from the MinHash sketch is 𝒪(*s*). Hence, the time complexity for calculating the pairwise distance matrix between all pairs of sequences from the sketches is 𝒪(*sn*^2^). For suitable applications of this algorithm however, both the sketch size and the number of sequences are numerically dominated by the combined length of the sequences. Thus, the expected time to run the algorithm is linear with respect to the combined length of the sequences. This has been verified empirically with the 960 REFSOIL dataset (Figure 9). The figure shows that as the number of sequences grows (and hence the combined length of the sequences), the time taken to construct the cluster tree grows linearly. When applied to the full REFSOIL dataset using 10 cores on a MacBook Pro M2 Max, `ctree` took ∼3.5 minutes.

**Figure 9.**
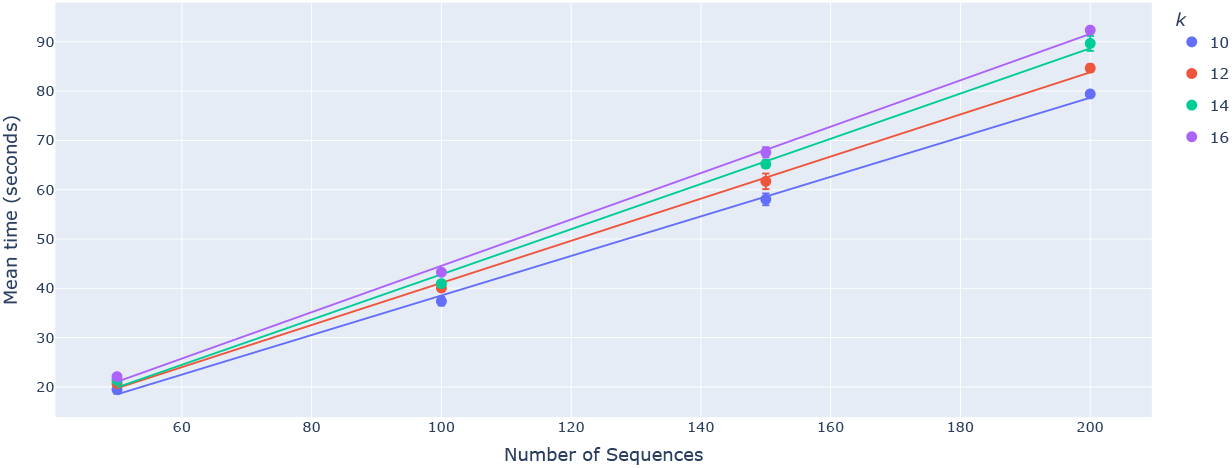
Computational performance of the `dvs_ctree` app on 960 REFSOIL microbial dataset using 8 cores. The wall time taken to run the algorithm grows linearly with respect to the number of sequences.

## Recommendations

For selecting representative sequences for large-scale analyses, we recommend use of the `nmost` command line tool. The choice of *k* should be guided by the maximum number of unique *k*-mers in a DNA sequence of length *L*, indicated as the result of the expression *log*(*L*/4). For instance, *k* ≈ 12 for bacterial genomes (which are of the order 10^6^bp). For individual protein coding genes, as Figure 4 indicates, *k* = 6 for `nmost` gives a reasonable accuracy. For estimating a phylogeny, we recommend *k* = 12 and a sketch size of 3000.

## Availability

`diverse-seq` can be installed from the Python Package Index. The GitHub repository for `diverse-seq` contains all the scripts used to generate the results in this work. A script for downloading the data sets used in this work, which are available from Zenodo **10.5281/zen-odo.14052787**, is also included.

## Acknowledgements

We thank Yapeng Lang for comments on the manuscript.

